# Immunological findings in a group of individuals who were non-responders to standard two-dose SARS-CoV-2 vaccines

**DOI:** 10.1101/2022.05.05.490815

**Authors:** Qiang Zeng, Xue Yang, Qi Gao, Biao-yang Lin, Yong-zhe Li, Gang Huang, Yang Xu

**Author notes:** These authors contributed equally to the study (Qiang Zeng, Xue Yang). **Correspondence:** Yang Xu, MD, PhD.

## Abstract

Coronavirus disease (COVID-19), caused by severe acute respiratory syndrome coronavirus 2 (SARS-CoV-2), was declared a pandemic. The virus has infected more than 505 million people and caused more than 6 million deaths. However, data on non-responders to SARS-CoV-2 vaccines in the general population are limited. The objective of the study is to comprehensively compare the immunological characteristics of non-responders to SARS-CoV-2 vaccines in the 18-59 years with that in the 60 years and older using internationally recognized cutoff values. Participants included 627 individuals who received physical examinations and volunteered to participate in COVID-19 vaccination from the general population. The main outcome was an effective seroconversion characterized by anti-SARS-CoV-2 spike IgG level of at least 4-fold increase from baseline. Profiling of naive immune cells was analyzed prior to vaccination to demonstrate baseline immunity. Outcomes of effective seroconversion in the 18-59 years with that in the 60 years and older were compared. The quantitative level of the anti-spike IgG was significantly lower in the 60 years and older and in men among the 18-59 years. There were 7.5% of non-responders among the 18-59 years and 11.7% of non-responders in the 60 years and older using the 4-fold increase parameter. The effective seroconversion rate was significantly related to the level of certain immune cells before vaccination, such as CD4 cells, CD8 cells and B cells and the age. An individual with a titer of anti-SARS-CoV-2 spike IgG that is below 50 BAU/mL might be considered a non-responder between 14-90 days after the last vaccine dose. Booster vaccination or additional protective measures should be recommended for non-responders as soon as possible to reduce disease severity and mortality.

---

Coronavirus disease (COVID-19), caused by severe acute respiratory syndrome coronavirus 2 (SARS-CoV-2), was declared a pandemic. The virus has infected more than 505 million people and caused more than 6 million deaths.^1^ Since December 2020 the World Health Organization (WHO) recommends vaccination against COVID-19, nine types of coronavirus disease 2019 (COVID-19) vaccines have been included in the emergency use list.^2^

Vaccination against COVID-19 is especially important in reducing severe illness and mortality. According to the data of the Centers for Disease Control and Prevention (CDC) in 2016–2017, the mortality rate caused by influenza virus was 0.13%.^3^

In order to bring the COVID-19 pandemic under control as soon as possible and ensure that the mortality rate of COVID-19 is close to that caused by influenza virus, the prevention and treatment of children as well as elderly and immunocompromised people has emerged as a top priority at present.^4–8^ Sun et al. have reported that hospitalization and severe outcomes were similar in unvaccinated healthy individuals and immunocompromised patients who received full two doses of SARS-CoV-2 vaccination in the United States, suggesting that COVID-19 breakthrough infection after SARS-CoV-2 vaccination is associated with immune dysfunction. Hospitalization and severe outcomes were 21.1% and 1.9%, respectively, in unvaccinated healthy individuals, and 20.7% and 2.1%, respectively, in patients with immune dysfunction after 14 days following full vaccination, indicating that an immune barrier is not well established in immunocompromised patients after full vaccination and post-vaccination serologic testing (PVST) is necessary to identify immunocompromised individuals without specific immunity so they can be given additional prophylaxis after full vaccination.^8^ This study suggests that PVST will help reduce mortality, showing the importance and urgency of PVST using an international standard.

To date, more than 5 billion people have been vaccinated against COVID-19.^9^ In clinical trials associated with COVID-19 vaccination, the effective COVID-19 vaccination reportedly elicits specific antibody responses. An effective humoral immune response is defined as a ≥ 4-fold increase in antibody titers from baseline and is considered gold standard for assessing antibody protection in vaccinated recipients in clinical studies.^10–12^ In contrast, a non-responder is an individual who demonstrates no effective humoral immune response despite the completion of the suggested vaccination procedure.^13–14^

Non-responders to the vaccine have been described in the hepatitis B vaccine. Szmuness et al. have reported that 7.4% of immunized individuals fail to elicit detectable specific antibodies after two doses of hepatitis B vaccine, suggesting that there are non-responders in the population.^15^ Roome et al. have found that 11.9% of individuals with hepatitis B vaccine were no or inadequate levels of antibody, suggesting that PVST should be done at intervals of 30 to 90 days after the last vaccine dose.^16^

Repeated non-responders to a third or fourth dose of SARS-CoV-2 vaccines were first observed in transplant recipients. Caillard et al. reported a cohort of 92 renal transplant recipients who did not have effective seroconversion after the third dose of mRNA vaccines.^17^ Furthermore, there were 52.9% (18/34) of non-responders to BNT162b2 vaccine and 48.3% (28/58) of non-responders to mRNA-1273 vaccine after the fourth dose of mRNA vaccines, showing that PVST should be done after the a third or fourth vaccine dose.

Non-responders to the SARS-CoV-2 vaccine are a vulnerable population with poor outcomes and high mortality. Chukwu et al. reported clinical findings in a group of kidney transplant recipients received two doses of the SARS-CoV-2 vaccine.^18^ There were 22 breakthrough infections and 3 deaths after vaccination, including 77% (17/22) infections and 13.6% (3/22) deaths in the non-responder group and only 23% (5/22) infections and 0% (0/22) deaths in the responder group.^18^ Therefore, there is an urgent need to identify SARS-CoV-2 vaccine non-responders in the vulnerable population to reduce severe COVID-19 and mortality.

During the promotion of vaccination, several factors affecting the response to the SARS-CoV-2 vaccine were taken into consideration, especially the reduced response to the SARS-CoV-2 vaccine in children, elderly people, and immunocompromised population. It has been documented that there are 5–10% of non-responders to hepatitis B vaccines in healthy individuals.^19^ However, the clinical characteristics of non-responders to SARS-CoV-2 vaccines in clinical trials remain limited.^20–24^

To this end, in December 2020, WHO issued an International Standard (IS) for the quantification of anti-SARS-CoV-2 immunoglobulin for PVST.^25–26^ This standard provides a unified benchmark for effective antibody protective concentrations after vaccination.

Zhang et al. reported a cohort of 75 healthy individuals who received two doses of the inactivated SARS-CoV-2 vaccine in 29-49 yr.^27^ There were 9.3% (7/75) of non-responders to the SARS-CoV-2 vaccine in the cohort study and a low lymphocyte count was a risk factor for non-responders. However, internationally recognized cutoffs according to the WHO IS and data from the elderly (≥60 years) were not reported.

The objective of this clinical study was to describe the immunological characteristics of 627 people in 18-86 yr that volunteered to participate in COVID-19 vaccination and to comprehensively compare the non-responders in the 18-59 yr population with the ≥60 yr population to explore internationally recognized cutoffs. Profiling of naïve immune cells were analyzed prior to vaccination to demonstrate baseline immunity. After two doses of vaccination, the antibody titer was evidently increased by ≥ 4 times from baseline as the gold standard. Furthermore, the data using the WHO IS cutoff was analyzed to provide insights for identifying non-responders and improving the efficacy of vaccines, help in reduction of breakthrough infections after vaccines, and ultimately to reduce the disease severity and mortality.

## Results

### Baseline immunological characteristics of individuals prior to vaccination

There were 627 individuals with anti-RBD/Spike IgM and IgG negative prior to vaccination, and 42.4% (266/627) individuals aged ≥60 yr and 50.9% (319/627) male enrolled in the study (Table 1). In the 18-59 yr group, the medians [interquartile ranges (IQRs)] of absolute lymphocyte count (ALC), CD4 cell count, CD8 cell count, B cell count, and natural killer (NK) cell count were 1,476 (1,168–1,875), 851 (677–1,151), 490 (357–632), 256 (179–367), and 193 (141–287)/mm^3^, respectively. On the contrary, in the ≥60 yr group, the respective medians (IQRs) were 1,281 (1,023–1,520), 747 (562–955), 418 (288–544), 204 (138–303), and 234 (162–355)/mm^3^ (Table 1). In fact, the number of naïve lymphocytes, CD4 cells, CD8 cells, and B cells were significantly reduced in the elderly (≥60 years) population than that in the 18–59 yr population (*P* < 0.001). Hence, these naïve immune cells wane significantly, while the NK cell counts increase significantly in the elderly people (Table 1, Figure 1A).

**Figure 1.**
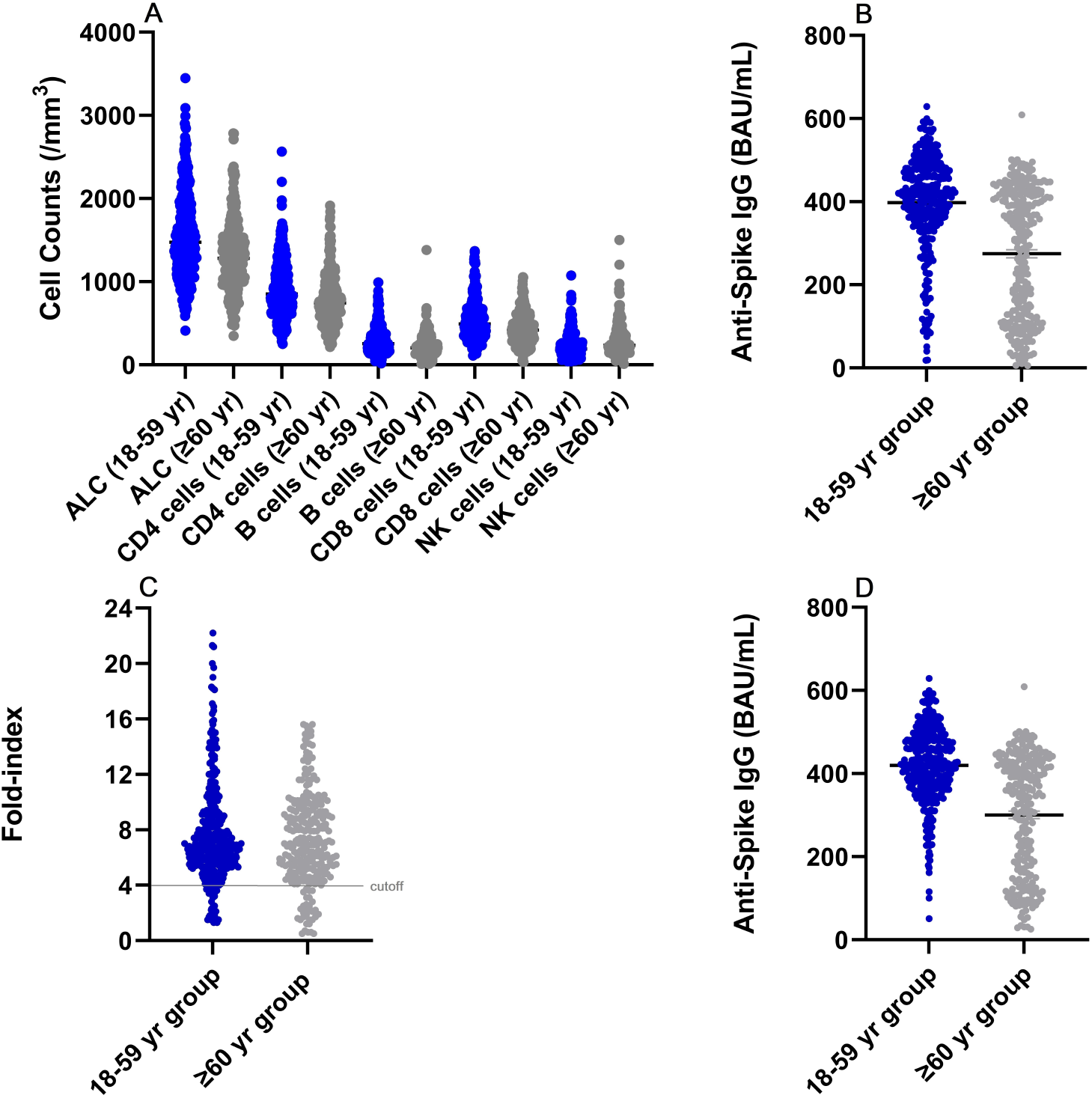
Immunological characteristics of 627 individuals. (A) Naïve cellular immunological parameters of the 627 cases who received physical examinations. These naïve immune cells wane significantly, while the natural killer (NK) cell counts increase significantly in the elderly people. ALC, absolute lymphocyte count. (B) The anti-spike IgG levels after complete vaccination of the 627 cases. The quantitative level of the anti-spike IgG was significantly lower in the ≥60 yr group (median 307.2, IQR 118.2–417.3 BAU/mL) than that in the 18-59 yr group (median 416.8, IQR 355.7–479.2 BAU/mL, *P* < 0.001. Mean and standard error of the mean (SEM) were shown. (C) The vaccine-induced responses using at least a 4-fold increase in antibody titer from baseline in 627 cases. There were 7.5% of non-responders (fold-index < 4) among the 18–59 yr group and 11.7% in the ≥60 yr group. The level of anti-spike IgG ranges (the 1st–99th percentile) for responders (fold-index ≥ 4) were 43.9–592.0 BAU/mL in combination of the 18–59 yr group and the ≥60 yr group. A cutoff line at fold-index 4 was shown. (D) In the responder group (fold-index ≥ 4), levels of anti-spike IgG for the 1st–99th percentile were 131.8-592.3 BAU/mL in the 18–59 yr group, and 29.7–500.9 BAU/mL in the ≥60 yr group, respectively. Mean and SEM were shown.

**Table 1.** Immunological characteristics of individuals before and after vaccination.

| Characteristics | 18–59 yr group | ≥60 yr group | *P* value |
| --- | --- | --- | --- |
| **Total number of cases** | 361 | 266 |  |
| **Sex (%)** |  |  |  |
| **Male** | 183 (50.7) | 136 (51.1) |  |
| **Female** | 178 (49.3) | 130 (48.9) |  |
| **Age, yrs mean (SD)** | 45 (9) | 67 (6) |  |
| **Naïve immune cells, median (IQR)** |  |  |  |
| **Lymphocytes (/mm<sup>3</sup>)** | 1,476<br>(1,168–1,875) | 1,281<br>(1,023–1,520) | <0.001 |
| **CD4 cells (/mm<sup>3</sup>)** | 851<br>(677–1,151) | 747<br>(562–955) | <0.001 |
| **CD8 cells (/mm<sup>3</sup>)** | 490<br>(357–632) | 418<br>(288–544) | <0.001 |
| **B cells (/mm<sup>3</sup>)** | 256<br>(179–367) | 204<br>(138–303) | <0.001 |
| **Natural killer cells (/mm<sup>3</sup>)** | 193<br>(141–287) | 234<br>(162–355) | <0.001 |
| **Anti-spike IgG** |  |  |  |
| **Seropositivity, % (no.)\*** | 99.7 (360/361) | 98.9 (263/266) |  |
| **Titer, BAU/mL, median (IQR)** | 416.8 (355.7–479.2) | 307.2 (118.2–417.3) | <0.001 |

\*The cutoff value for seropositivity was 11.6 BAU/mL. SD, standard deviation; IQR, interquartile range; IgG, immunoglobulin G; CD, cluster of differentiation

### Characteristics of humoral response after complete vaccination

We analyzed the anti-spike immunoglobulin (Ig)G levels after complete two doses of vaccination of the 627 cases (Table 1). Post-vaccination testing was done at intervals of 14 to 90 days after the second vaccine dose. The anti-spike IgG seropositive rates were 99.7% in the 18–59 yr population and 98.9% in the ≥60 yr population based on the cutoff (the cutoff value for seropositivity was 11.6 BAU/mL) (Table 1). However, the quantitative level of the anti-spike IgG was significantly lower in the ≥60 yr group (median 307.2, IQR 118.2–417.3 BAU/mL) than that in the 18–59 yr group (median 416.8, IQR 355.7–479.2 BAU/mL) (Table 1, Figure 1B). The reference ranges (the 2.5th–97.5th percentile) were 88.9–576.2 BAU/mL in the 18–59 yr group and 27.7–491.0 BAU/mL in the ≥60 yr group at intervals of 14–90 days after complete vaccination (Table 2).

**Table 2.** Characteristics of immunity before and after the complete vaccination.

| Characteristics | 18–59 yr group |  |  | ≥60 yr group |  |  |
| --- | --- | --- | --- | --- | --- | --- |
|  | Fold-index <4 | Fold-index ≥4 | *P* value | Fold-index <4 | Fold-index ≥4 | *P* value |
| **Total number of cases** | 361 |  |  | 266 |  |  |
| **Anti-spike IgG, BAU/mL**<br>**(the 2.5th–97.5th percentile)** | 88.9–576.2 |  |  | 27.7–491.0 |  |  |
| **Fold-index, % (no.)\*** | 7.5 (27/361) | 92.5 (334/361) |  | 11.7 (31/266) | 88.3 (235/266) |  |
| **Anti-spike IgG, BAU/mL** |  |  |  |  |  |  |
| **Median (IQR)** | 115.8 (88.6–167.8) | 420.8 (369.9–480.6) | <0.0001 | 63.9 (35.1–106.9) | 346.0 (160.4–424.7) | <0.0001 |
| **The 2.5th–97.5th percentile** | 11.3–266.3 | 200.7–576.5 |  | 5.4–317.8 | 46.6–491.1 |  |
| **Naïve immune cells (/mm<sup>3</sup>)** |  |  |  |  |  |  |
| **Lymphocytes, mean (95% CI)** | 1,130 (1,007–1,252) | 1,578 (1,524–1,633) | <0.0001 | 1,015 (888–1,143) | 1,344 (1,291–1,397) | <0.0001 |
| **CD4 cells, mean (95% CI)** | 631 (555–708) | 942 (905–979) | <0.0001 | 563 (494–631) | 818 (777–858) | <0.0001 |
| **CD8 cells, mean (95% CI)** | 414 (349–479) | 532 (508–557) | 0.0081 | 394 (310–478) | 444 (420–468) | 0.1744 |
| **B cells, mean (95% CI)** | 119 (72–166) | 306 (289–323) | <0.0001 | 74 (60–88) | 248 (231–266) | <0.0001 |
| **NK cells, mean (95% CI)** | 192 (151–233) | 235 (220–251) | 0.1241 | 281 (225–337) | 286 (261–311) | 0.8902 |

\*Post-vaccination testing was performed at intervals of 14–90 days after the second vaccine dose. The reference ranges were defined as the 2.5th–97.5th percentile in the study. NK, natural killer; IQR, interquartile range; IgG, immunoglobulin G; CD, cluster of differentiation; CI, confidence interval; SARS-CoV-2, severe acute respiratory syndrome coronavirus 2

### Characteristics of humoral response in male and female after complete vaccination

The quantitative level of the anti-spike IgG was significantly lower in the male group (median 404.9, IQR 326.7–471.7 BAU/mL) than that in the female group (median 421.7, IQR 367.1– 480.7 BAU/mL) in 18–59 yr group (*P* = 0.0008) (Figure 2A). There was no significant difference in the ≥60 yr group for the quantitative levels of anti-spike IgG between the male group (median 285.4, IQR 113.1–416.3 BAU/mL) and the female group (median 327.5, IQR 126.7–418.2 BAU/mL) (*P* = 0.4517) (Figure 2B).

**Figure 2.**
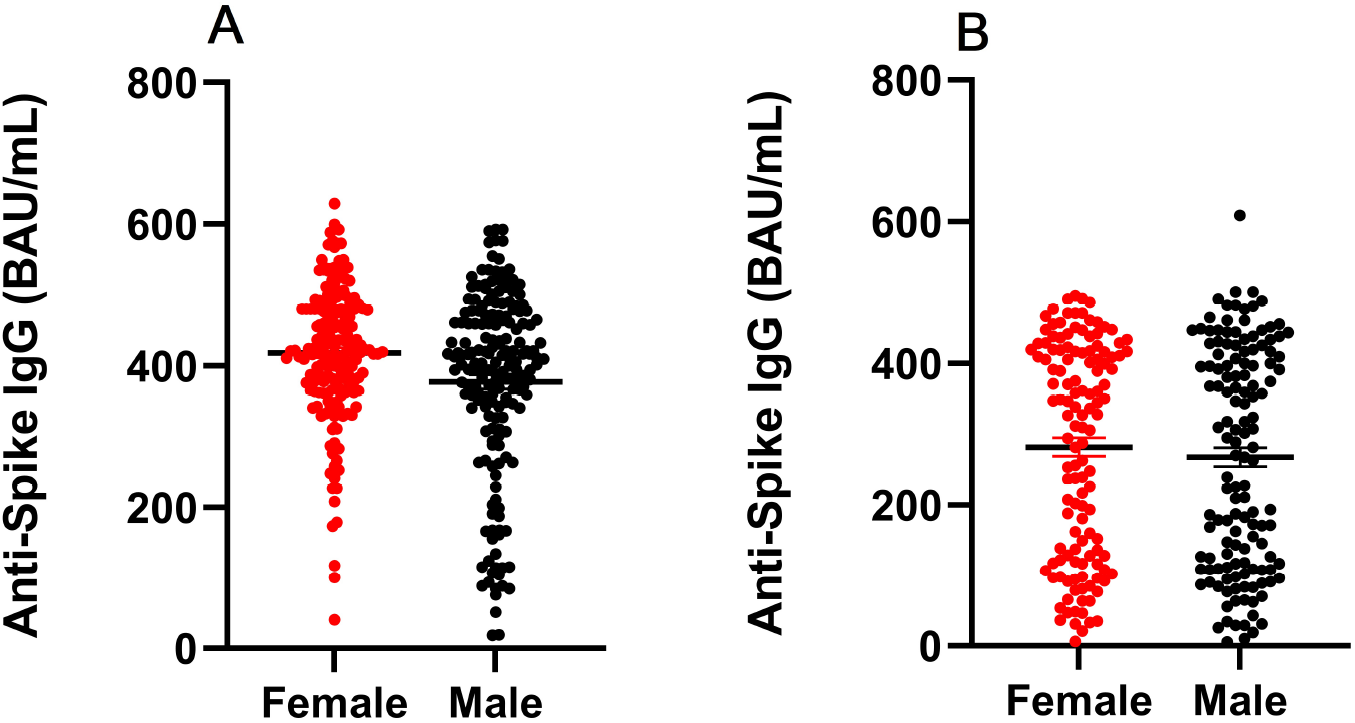
Sex affected anti-spike IgG levels after complete vaccination. (A) The quantitative level of the anti-spike IgG was significantly lower in the male group (median 404.9, IQR 326.7–471.7 BAU/mL) than that in the female group (median 421.7, IQR 367.1–480.7 BAU/mL) in 18–59 yr group (*P* = 0.0008). Mean and SEM were shown. (B) There was no significant difference in the ≥60 yr group for the quantitative levels of anti-spike IgG between the male group (median 285.4, IQR 113.1–416.3 BAU/mL) and the female group (median 327.5, IQR 126.7–418.2 BAU/mL) (*P* = 0.4517). Mean and SEM were shown. IQR, interquartile range; SEM, standard error of the mean.

### Characteristics of responders and non-responders to the SARS-CoV-2 vaccine using internationally recognized standards

Thereafter, we evaluated the vaccine-induced responses, based on the post-second-dose and pre-second-dose titers, using the 4-fold increase parameter (fold-index <4 or ≥4) and the WHO IS (Table 2). Remarkably, there were 7.5% of non-responders (fold-index < 4) among the 18–59 yr group and 11.7% of non-responders in the ≥60 yr group (Table 2, Figure 1C), indicating that the positive rate of anti-spike IgG cannot represent the effective seroconversion rate (fold–index ≥ 4). Therefore, the anti-spike IgG positivity or seroprevalence might not be a suitable predictor of the effective seroconversion.

In the 18–59 yr group, the median (IQR) levels of anti-spike IgG and the reference ranges were 115.8 (88.6–167.8) and 11.3-266.3 BAU/mL with fold-index < 4, respectively and 420.8 (369.9–480.6) and 200.7-576.5 BAU/mL with fold-index ≥ 4, respectively (*P*< 0.0001) (Table 2). In contrast, in the ≥60 yr group, the median (IQR) levels of anti-spike IgG and the reference ranges were 63.9 (35.1–106.9) and 5.4-317.8 BAU/mL with fold-index < 4, respectively and 346.0 (160.4–424.7) and 46.6-491.1 BAU/mL with fold-index ≥ 4, respectively (*P* < 0.0001) (Table 2). The level of anti-spike IgG ranges (the 1st-99th percentile) for all responders (fold-index ≥ 4) were 43.9-592.0 BAU/mL in combination of the 18-59 yr group and the ≥60 yr group at 14-90 days after complete vaccination (Figure 1C).

### Relationship between effective seroconversion rate and baseline immunity

We further observed that the effective seroconversion rate (fold–index ≥ 4) was significantly related to the level of certain naïve immune cells before vaccination (Table 2). Particularly, the lymphocyte count was significantly different (*P* < 0.0001) between the fold-index <4 or ≥4 groups. For instance, in the 18–59 yr group, the lymphocyte count was 1,130/mm^3^ [95% CI (1,007–1,252)/mm^3^] in the <4 group and 1,578/mm^3^ [95% CI (1,524–1,633)/mm^3^] in the ≥4 group. On the contrary, in the ≥60 yr group, the lymphocyte count was 1,015/mm^3^ [95% CI (888–1,143)/mm^3^] in the <4 group and 1,344/mm^3^ [95% CI 1,291-1,397)/mm^3^] in the ≥4 group. Similarly, the CD4 cell counts were significantly different (*P* < 0.0001) between the individuals with fold-index <4 and ≥4. For instance, in the 18–59 yr population, the CD4 cell count was 631/mm^3^ [95% CI (555–708)/mm^3^] versus 942/mm^3^ [95% CI (905–979)/mm^3^] in the <4 and ≥4 groups, respectively, while in the ≥60 yr age group, it was 563/mm^3^ [95% CI (494–631)/mm^3^] versus 818/mm^3^ [95% CI (777–858)/mm^3^] in the <4 and ≥4 groups, respectively. With respect to the B cell count, there was a significant difference (*P* < 0.0001) between the individuals with fold-index < 4 and ≥ 4. For example, in the 18-59 age group, the B cell count was 119/mm^3^[95% CI (72–166)/mm^3^ versus 306/mm^3^ [95% CI (289–323)/mm^3^] in the <4 and ≥4 groups, respectively, whereas in the ≥60 yr age group, it was 74/mm^3^ [95% CI (60-88)/mm^3^] versus 248/mm^3^ [95% CI (231–266)/mm^3^] in the <4 and ≥4 groups, respectively. Regarding the CD8 cell count, a significant difference was noted only between the individuals with <4 and ≥4 fold-indices in the 18–59 yr age group [414/mm^3^ (95% CI 349–479/mm^3^) versus 532/mm^3^ (95% CI 508–557/mm^3^), *P*= 0.0081]. However, the CD8 cell count in the ≥60 yr group and NK cell count in both the age groups did not portray any significant differences.

## Discussion

To the best of our knowledge, this is one of the first clinical studies to report non-responders after administering the full two doses of inactivated SARS-CoV-2 vaccines using the WHO International Standard (IS) cutoff value for anti-SARS-CoV-2 immunoglobulin (Ig) with documented baseline immunity & effective seroconversion using antibody titers ≥ 4 times from baseline as the gold standard. Previous clinical trials have demonstrated that individuals have similar neutralizing capacity against the D614G and B.1.1.7 variants compared with wild-type pseudovirus after received two doses of the SARS-CoV-2 HB02 vaccine through cross-reactivity^24^.

Whether there is a humoral immune response following COVID-19 vaccination is a marker of population immunity.^28–30^ Typically, health individuals with normal immune systems have normal immune cell counts and an effective humoral immune response, defined as a ≥4-fold rise in antibody titers from baseline within 14-90 days of the vaccination schedule. The use of anti-SARS-CoV-2 assays with the WHO IS can facilitate the comparison of the strength of the humoral immune response between individuals, making the data more accurate and providing reliable data for the COVID-19 vaccine booster. Therefore, adequate clinical trials are necessary regarding the assessment of immune characteristics of individuals prior to vaccine booster shot, such that the mortality in the pandemic may be quickly reduced.

We used an anti-SARS-CoV-2 spike quantitative IgG kit (COVID-SeroKlir Kantaro SARS-CoV-2 IgG Ab Kit) approved by the Food and Drug Administration (FDA) under Emergency Use Authorization (EUA) with the WHO IS. This kit has been extensively evaluated in many clinical studies, including neutralizing antibodies after SARS-CoV-2 infection, immunological memory to SARS-CoV-2, convalescent plasma treatment of severe COVID-19, and antibody responses to mRNA vaccines in healthy people and patients.^30–35^ After complete two dose vaccination, the level of anti-spike IgG ranges (the 1st–99th percentile) for all responders (fold–index ≥ 4) were 43.9–592.0 BAU/mL. A preliminary cutoff of 50 BAU/mL was set based on percentiles of all responders and convenience of manufacturing standard controls. The final cutoff value will be determined by future clinical trials.

Both our and Zhang’s data showed that approximately 10% of the studied population did not respond to the inactivated SARS-CoV-2 vaccine following full two doses, indicating the importance of monitoring non-responders in this population.^27^ Similar to Zhang’s report, sex affecting anti-spike IgG levels among the 18–59 yr group after complete vaccination was also observed in our study due to female with higher estradiol hormone.^27^ However, Zhang et al. did not report data on the elderly. Our data showed that there was no significant difference in the quantitative levels of anti-spiking IgG between the male and female groups aged ≥60 years (P = 0.4517), suggesting that female estradiol hormone declines over the age of 60 years.

The WHO IS has demonstrated to be enabled to comparison between different types of vaccines. Zitt et al. reported that the median titers of non-seroconversion and seroconversion were 635.5 and 1,565.0 BAU/mL after two doses of mRNA vaccination in hemodialysis patients at 67.6 ± 14.8 years, respectively;^36^ whereas we reported that the median titers of non-seroconversion and seroconversion were 63.9 and 346.0 BAU/mL after giving two doses of inactivated SARS-CoV-2 vaccines at 67 ± 6 years, respectively, indicating that the mRNA vaccines is more potent than the inactivated SARS-CoV-2 vaccines.^37^

The benefits of post-vaccination serologic testing (PVST) outweigh the potential risks. Zitt et al. reported there were median titer of 1,440 BAU/mL in documented hepatitis B virus (HBV) vaccine responders (anti-HBs antibody ≥10 mIU/mL) and median titer of 308.5 BAU/mL in non-responders after two doses of mRNA vaccination (*P* = 0.035), suggesting that PVST might predict the general immune competence.^36^ All anti-SARS-CoV-2 spike IgG-positive patients recovered from the infection respond well to the vaccine, which indirectly proves this phenomenon.^29–30^ If this theory turns out to be correct, then it is possible that SARS-CoV-2 vaccine responders with normal immune cell counts have a strong ability to produce antibodies against variants through asymptomatic infections or cross-reactivity.^24^ This may support the Government-issued “immunity passports” to demonstrate an individual’s immune ability according to the WHO IS (≥ 50 BAU/mL) after recovered from COVID-19 or following the SARS-CoV-2 vaccine.

The most significant benefit of PVST is to save patient lives. There were 2.1% of severe outcomes for COVID-19 breakthrough infection in immunocompromised patients after two vaccine doses.^8^ However, we can identify these with anti-spike IgG below 50 BAU/mL to reduce mortality. Chukwu et al. reported clinical findings in a group of kidney transplant recipients (KTRs) received with two doses of vaccines (72% of BNT162b2, 28% of AZD1222). There were 22 breakthrough infections and 3 deaths after vaccination, including 77% (17/22) infections and 13.6% (3/22) deaths in the seronegative group and only 23% (5/22) infections and 0% (0/22) deaths in the seropositive group.^18^ Chavarot et al. reported that administration of a third dose of BNT162b2 vaccine did not improve immunogenicity in KTRs treated with belatacept without prior COVID-19. Seropositivity was only 37.1% (13/35) of KTRs after the third vaccine dose. Twelve KTRs developed symptomatic COVID-19 after vaccination, including severe outcomes (50% of mortality).^38^ However, those studies did not use the WHO IS to get a cutoff for the responder. Therefore, there may be some non-responders (fold-index <4) in the seropositive group according to our study.

For SARS-CoV-2 vaccine non-responders, one benefit from PVST to the patient is to get a booster shot as soon as possible.^39–40^ For persistent non-responders to SARS-CoV-2 vaccination, anti-SARS-CoV-2 immunoglobulin injections could save these lives in the seronegative group following the vaccination.^41–43^

Another good example for PVST is the hepatitis B vaccine. After the first hepatitis B vaccine was approved in the United States in 1981 and the recombinant hepatitis B vaccine developed by Maurice Hilleman was approved by the FDA in 1986, it took scientists many years to realize that the vaccine did not provide good protection for the elderly and certain immunocompromised populations and put them at risk of breakthrough infections after vaccination.^44–45^ Szmuness et al. have first reported that 7.4% of immunized individuals fail to elicit detectable specific antibodies after two doses of hepatitis B vaccine, suggesting that there are non-responders in the population.^15^ Roome et al. have found that 11.9% of individuals with hepatitis B vaccine were no or inadequate levels of antibody, suggesting that PVST should be done at intervals of 30 to 90 days after the last vaccine dose.^16^ Non-responders to hepatitis B vaccine were observed in adults, infants as well as in drug users.^13, 46,47^ Many subsequent studies have shown that the elderly and immunocompromised populations are associated with reduced vaccine responses to hepatitis B vaccination.^45–47^ The CDC has recommended PVST using the WHO IS for immunocompromised individuals following hepatitis B vaccine based on evidence of non-responders in the population.^45^ For persistent non-responders (anti-HBs antibody <10 mIU/mL, the WHO IS 07/164) to hepatitis B vaccination, anti-hepatitis B immunoglobulin injections are recommended if exposed to the hepatitis B virus.^45^

Furthermore, the lower baseline immunity may form the major risk factor for non-responders after administration of full SARS-CoV-2 vaccine in our study. Van Oekelen et al. have demonstrated that 32.3% (10/31) of multiple myeloma patients with severe lymphopenia (<500/mm^3^) remained negative for SARS-CoV-2 spike IgG after two doses of mRNA vaccines (OR 2.89, 95% CI 1.10-7.20, *P* = 0.018).^48^ Similarly, two studies reported that 63.7-77.3% of patients who had a history of anti-CD20 therapy for B cell depletion remained negative for SARS-CoV-2 IgG after receiving mRNA vaccines, suggesting that B cells are required for humoral immunity following COVID-19 vaccines.^49–50^ Hence, further clinical trials must be performed to finalize effective booster shots for immunocompromised people after administering the complete dose in the general population.^51–55^

Existing data show that laboratory testing has certain guiding significance for the prevention and treatment of COVID-19: (1) a normal immune cell count and a good response to the SARS-CoV-2 vaccine indicate healthy individuals; (2) a decrease in immune cell count may predict the disease severity and severe outcomes;^6,56^ (3) anti-SARS-CoV-2 spike IgG levels < 50 BAU/mL following the vaccine designate non-response to the SARS-CoV-2 vaccine. According to this study, the anti-spike IgG seropositivities were 99.7% and 98.9% in the 18-59 yr and ≥60 yr groups, respectively. Additionally, certain naïve immune cells, such as CD4 cells, CD8 cells, and B cells exhibited significant waning in the elderly people, suggesting that the non-seroconversion rates were higher in individuals with lower baseline immunity. Incidentally, the anti-spike IgG seroprevalence or positivity was inconsistent with effective seroconversion rates observed in our study, thereby suggesting that anti-spike IgG positivity might not be a suitable predictor for the effective seroconversion rates. Our data showed that 7.5–11.7% of non-responders existed in the population, even some non-responders with anti-spike IgG positivity, supporting the CDC’s concern that some non-responders are positive for anti-spike IgG after vaccination.^57^ An FDA EUA quantitative assay with the WHO IS (20/136) cutoff might help to address this issue.^25–26^

To sum up, approximately 10% of the studied population has no response to both hepatitis B vaccine and SARS-CoV-2 vaccine, which is the content of urgent research to identify non-responders in the population. At present, among the people who have received two doses of the SARS-CoV-2 vaccine in the world, there are still many people who have not received the third dose. We need to identify non-responders using PVST as soon as possible because they are susceptible and we need to give them the third dose in priority. If PVST is not performed, these non-responders will become a vulnerable population and easily develop severe COVID-19. However, there are several potential strategies that can be employed to reduce the COVID-19 mortality rate below 0.13 %of that caused by influenza virus. These include the following measures: (1) Increase the vaccination rate of the general population;^2^ (2) Develop vaccines and/or anti-SARS-CoV-2 immunoglobulins against emerging and potential variants;^58–62^ (3) Administer booster vaccines for non-responders;^63–64^ (4) Accelerate clinical trials of intranasal SARS-CoV-2 vaccines to prevent transmission;^65^ (5) Assessment of humoral immune response of children, the elderly, and immunocompromised persons within 1–3 months after vaccine booster shot;^45, 66–71^ and (6) Incorporate additional protective measures for individuals with persistent (a fourth or fifth dose) negative humoral immune response after booster vaccination, such as injection of anti-SARS-CoV-2 immunoglobulins, antiviral drug treatment, usage of N95 masks in endemic areas, etc.masks in endemic areas, etc.masks in endemic areas, etc.masks in endemic areas, etc.^41–43, 72–77^

## Conclusions

In this prospective clinical study, naïve immune cells such as CD4 cells, CD8 cells, and B cells and anti-spike IgG levels were significantly reduced in the elderly. There were 7.5% of non-responders to SARS-CoV-2 vaccines in the 18-59 yr group and 11.7% of non-responders in the ≥60 yr group. The effective seroconversion rate was significantly related to the level of certain naïve immune cells before vaccination, such as CD4 cells, CD8 cells and B cells and the age. Sex affected anti-spike IgG levels in the 18-59 yr group after standard two-dose SARS-CoV-2 vaccines. An individual with a titer of anti-SARS-CoV-2 spike IgG that is below 50 BAU/mL might be considered a non-responder between 14-90 days after the last vaccine dose.

### Limitations

Study limitations include small sample size, data was only sourced from a single center, and lack of RBD/spike-specific cellular immune assessments.

